# Precision combination therapies based on recurrent oncogenic co-alterations

**DOI:** 10.1101/2020.06.03.132514

**Authors:** Xubin Li, Elisabeth K. Dowling, Gonghong Yan, Behnaz Bozorgui, Parisa Imarinad, Jacob H. Elnaggar, Augustin Luna, David G. Menter, Scott Kopetz, Chris Sander, Anil Korkut

## Abstract

Cancer cells depend on multiple driver alterations whose oncogenic effects can be suppressed by drug combinations. Discovery of effective combination therapies is challenging due to the complexity of the biomolecular landscape of drug responses. Here, we developed the method REFLECT (REcurrent Features Leveraged for Combination Therapies), which integrates machine learning and cancer informatics algorithms. The method maps recurrent co-alteration signatures from multi-omic data across patient cohorts to combination therapies. Using the REFLECT framework, we generated a precision therapy resource matching 2,201 drug combinations to co-alteration signatures across 201 cohorts stratified from 10,392 patients and 33 cancer types. We validated that REFLECT-predicted combinations introduce significantly higher therapeutic benefit through analysis of independent data from comprehensive drug screens. In patient cohorts with immunotherapy response markers, HER2 activation and DNA repair aberrations, we identified therapeutically actionable co-alteration signatures shared across patient sub-cohorts. REFLECT provides a framework to design combination therapies tailored to patient cohorts in data-driven clinical trials.

## Introduction

Targeted therapies such as HER2 and PARP inhibitors have significantly improved clinical outcomes in cancer^1,2,3^. Despite the transformative advances in sequencing technologies and precision medicine, durable responses to genetically-matched targeted therapies have remained limited^4,5^. The limited success can be at least partially attributed to treatment with mono-therapies targeting individual oncogenic aberrations. Oncogenesis, however, is driven by co-occurring aberrations in DNA, mRNA and protein levels that lead to co-activated oncogenic processes. A rich repertoire of preclinical and clinical studies has demonstrated that rational combination therapies blocking multiple oncogenic processes are more effective than mono-therapies^6,7,8^. Discovery of combination therapies that may benefit large patient cohorts, however, is a daunting task due to the complexity of the actionable oncogenic aberration landscapes^9^.

There is growing evidence that co-targeting the co-occurring oncogenic driver aberrations may induce effective and durable therapeutic responses. For example, PI3K-AKT pathway co-alterations alongside BRAFV600E mutations impact response to BRAF inhibitors and establish a rationale for co-targeting the BRAF and PI3K-AKT pathways^10^. Similarly, co-alterations in KRAS, BRAF and PIK3CA modulate response to EGFR/HER2 targeted therapies in colorectal cancers^8^. Genomic studies focusing on co-driver events in KRAS mutated lung cancers^11^ and recent combination therapy trials focusing on co-alterations (I-PREDICT^27^) also strongly support this argument^6,8^. Bioinformatics methods and resources that extract clinically useful knowledge (e.g., recurrent, co-occurring, co-actionable oncogenic events) from large multi-omic (genomic, transcriptomic and proteomic) data sets will accelerate the implementation of such precision combination therapies.

Here, we developed the machine learning algorithm and precision oncology resource termed “Recurrent Features Leveraged for Combination Therapy” (REFLECT). First, REFLECT generates multi-omic (DNA, mRNA, phosphoprotein) alteration signatures within a patient cohort (e.g., EGFR mutated patients) using a LASSO-penalty based sparse hierarchical clustering scheme^12^ followed by a statistical evaluation of significantly recurrent co-alterations. Each signature is marked by the distribution of recurrent co-alterations that can discriminate patients into sub-cohorts. Next, REFLECT maps the recurrent and co-actionable oncogenic alterations within multi-omic signatures of sub-cohorts to potential combination therapies based on the precision oncology knowledge bases. In the current analysis, each patient cohort is constructed based on the existence of at least one potentially actionable aberration with sampling from a pan-cancer patient meta-cohort (N=10392, TCGA resource)^13^. We have determined comprehensive genomic, transcriptomic and proteomic REFLECT-signatures as well as combination therapies matching to patient sub-cohorts in 201 unique cohorts. Through REFLECT analysis of the patient cohorts and proof-of-concept experimental validations in molecularly matched cell lines, we identified a series of drug combinations that can benefit patients marked by clinically relevant events such as PDL1 expression/immune infiltration, DNA checkpoint/repair aberrations and increased HER2 activity. We provide a searchable online resource with interactive visualization tools and tutorials (https://bioinformatics.mdanderson.org/reflect/). We also provide a REFLECT R package for analyses using custom patient data (https://github.com/korkutlab/reflect). We expect the tool and the associated resource will guide the discovery of combination therapies and selection of potentially benefiting patients for emerging combination therapy clinical trials.

## Results

### REFLECT-based signatures of oncogenic co-alterations

We generated a map of recurrent and co-actionable oncogenic alterations using the REFLECT pipeline (Figure 1). The analysis benefited from mutation, copy number, mRNA expression and phosphoproteomic analysis of 10,392 patients from 33 cancer types (TCGA) and 1601 cell lines from 34 lineages (CCLE, COSMIC cell lines project (COSMIC-CLP) and MDACC cell line project (MCLP)) (Figure 2, S1 and Table S1). To generate a meta-cohort with no batch effects, we used the protein and mRNA expression data from diverse cancer types (TCGA) that had been integrated and batch normalized with replica based normalization (RBN) and EB++ methods^14,15^. We also incorporated molecular profiling data from cell lines using a Z-score based normalization (see Methods). The integration of cell line data enables proof-of-principle experimental validations for matching drug combinations to specific signatures in molecularly matched lines. Next, we partitioned the cell line and patient pan-cancer meta-cohort into patient cohorts (stratified dataset), each defined by the existence of a “master” biomarker that is shared by all members of the cohort and potentially predictive of responses to a targeted cancer therapy (Figure 1B). The master biomarkers include potential therapeutically actionable genomic alterations such as those listed in the MDACC Precision Oncology Knowledge Base (N=182), and NCI-MATCH precision therapy trials (N=15)^16,17,18^. We have defined and added an immunotherapy cohort based on PDL1 overexpression and immune cell infiltration, two potential markers of immunotherapy response in diverse cancer types. Additional cohorts were constructed based on the existence of key oncogenic alterations, MYC amplifications, and TP53 and KRAS mutations. In total, we generated 201 biomarker-based cohorts. As provided in OncoKB resource, at least 25 of the “master” biomarkers are FDA-recognized and 14 biomarkers are NCCN-recommended as predictive of response to targeted therapy in clinical applications. The remaining biomarkers are under evaluation in precision oncology trials at MDACC and elsewhere.

**Figure 1.**
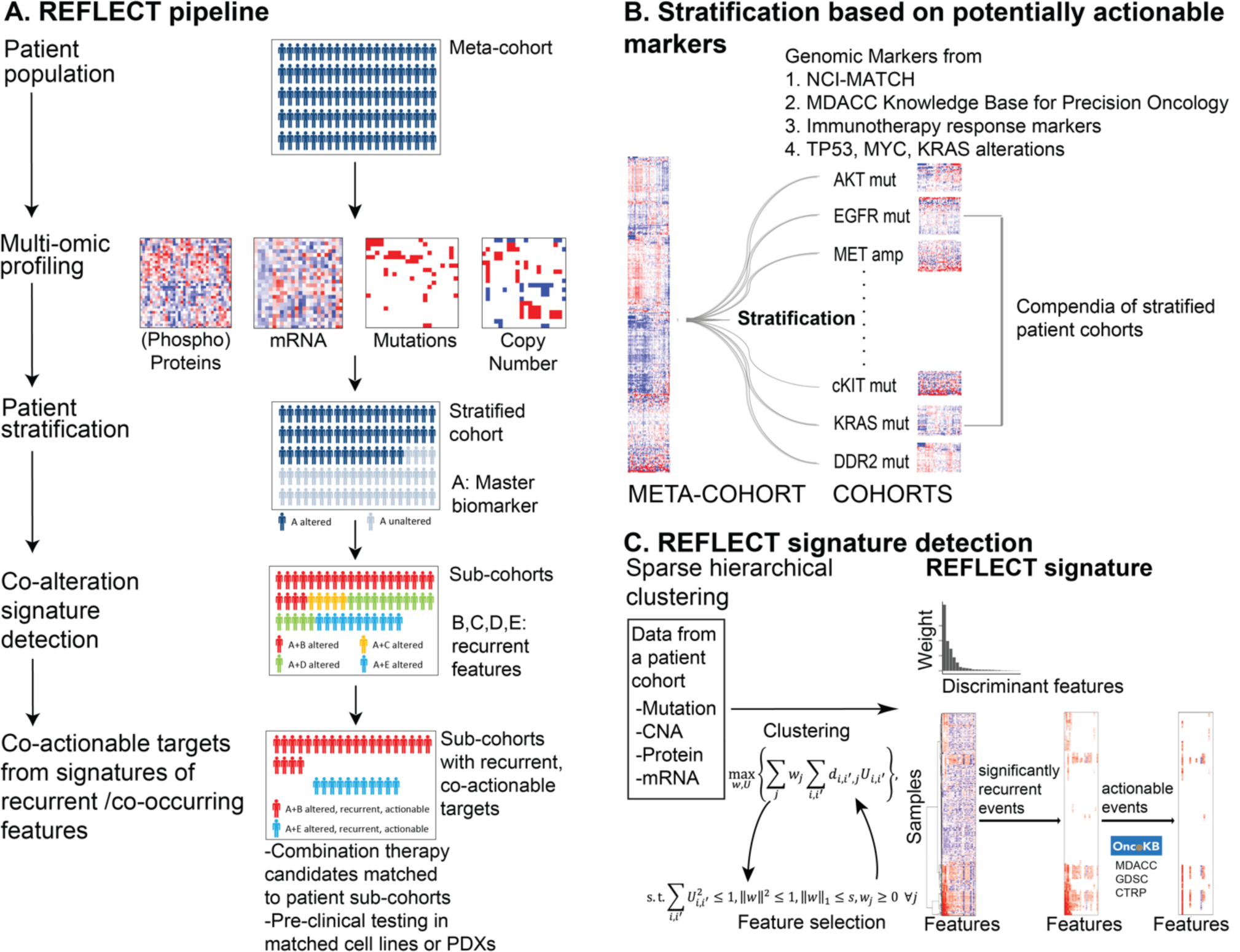
REFLECT. **A. The pipeline**. REFLECT identifies recurrent, co-occurring and co-actionable aberrations within patient cohorts. The method uses mutation, copy number, mRNA, and phosphoproteomics data. Patient cohort-specific multi-gene/protein signature detection with clustering/feature selection is followed by matching the signatures to combination therapies through statistical analyses of recurrence and automated searches in oncology knowledge bases. **B. Actionable alteration based cohort stratification**. The integrated pan-cancer data is partitioned into overlapping cohorts based on the existence of actionable genomic aberrations (e.g., cohort of EGFR mutant tumors). The mutational, copy number, mRNA and protein datasets from each partition are analyzed separately. **C. Selection of significantly recurrent alterations**. To identify sub-cohorts of patients characterized by a small set of discriminant and recurrent features (i.e., genomic alterations), the algorithm couples feature selection with hierarchical clustering. Once the recurrent discriminant features are detected, co-actionable targets within each sub-cohort (sub-cluster) are selected through statistical assessment of significantly recurrent patterns and matching features to drugs using precision oncology knowledge bases.

**Figure 2.**
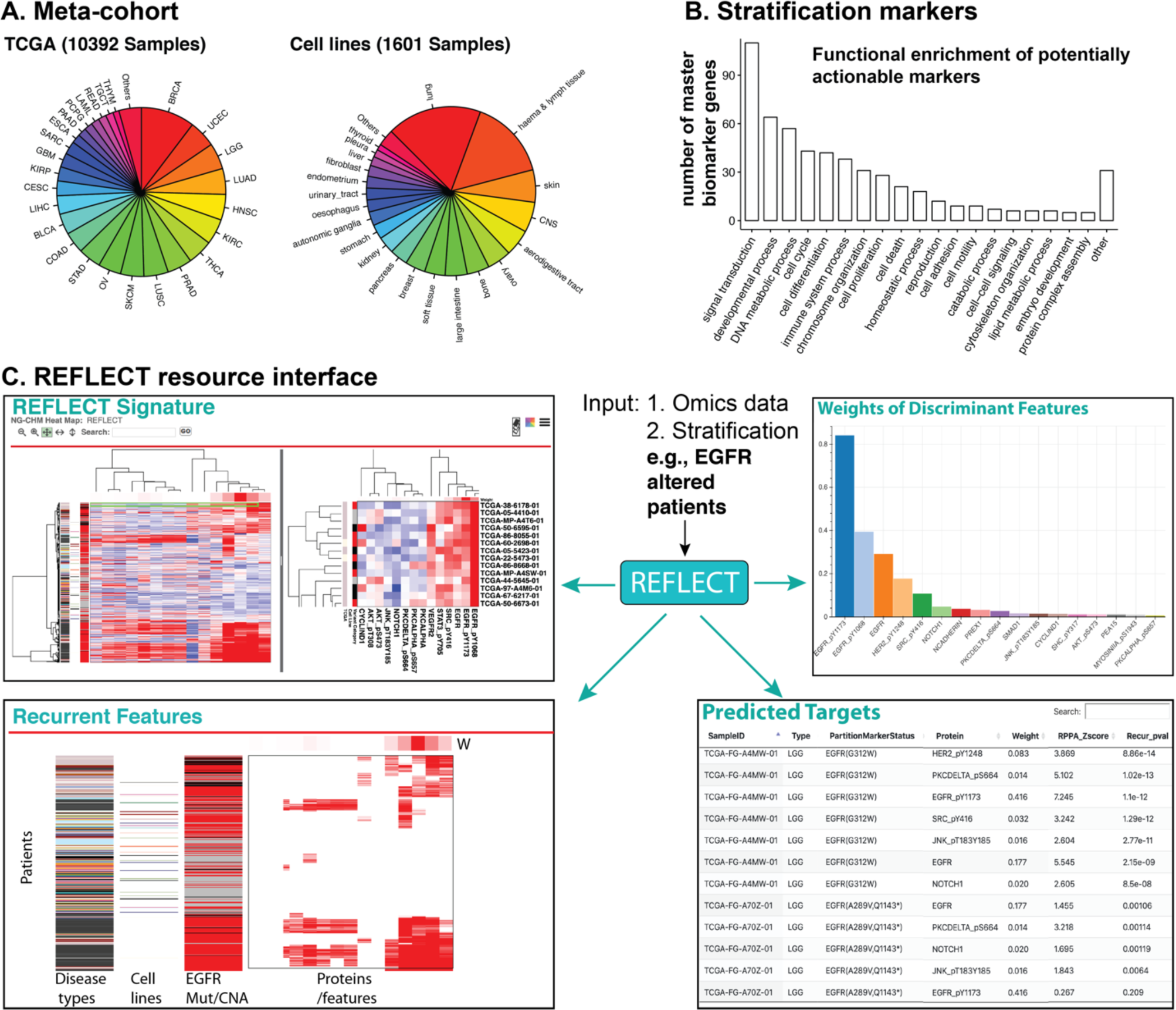
REFLECT resource and interface. **A. The REFLECT resource** benefits from a pan-cancer meta-cohort of cancer patient samples and cell lines from CCLE, COSMIC-CLP and MCLP. **B. The pan-cancer meta-cohort** is stratified into 201 sub-cohorts based on the existence of actionable master biomarkers. The master biomarkers are associated with diverse oncogenic processes as demonstrated with a functional GO-term enrichment analysis. **C. REFLECT resource outputs** the (i) distributions of disease types in each cohort, (ii) the most discriminant features (DNA, RNA, protein alterations) across each cohort, (iii) the REFLECT signature defined by the distribution of most discriminant features and (iv) selection of co-actionable targets based on the master biomarker and the REFLECT signature.

For each biomarker-based cohort, we generated the multi-omic/multi-gene signatures, termed as REFLECT-signatures, using the sparse hierarchical clustering algorithm^12^ (Figure 1C). The algorithm iterates LASSO-based feature detection and hierarchical clustering until convergence to a final signature (see methods). In succession of iterations, the feature detection selects a small set of alterations that contribute most to discrimination of patient sub-cohorts, which are identified by the clustering. A signature, which is shared by all members of the associated cohort, carries a combination of the master biomarker and a small set of REFLECT detected molecular features with non-zero contribution (***w***) to discrimination of patient sub-cohorts. To demonstrate the robustness of the REFLECT, we quantified the concordance between REFLECT signature features that were calculated with complete and subsampled partial data with varying coverage (60% to 95%) (Suppl Figure S4, S5). The concordance between partial and complete datasets increased monotonically with increased coverage such that concordance reaches an average of 83% for 70% coverage for the 8 patient cohorts tested (Figure S5). In summary, we have acquired a compendium of robust multi-omic REFLECT signatures spanning the patient cohorts identified by potentially co-actionable “master” biomarkers and sub-cohort specific features selected by our algorithm.

### REFLECT signatures inform the selection of recurrent and co-actionable targets

Next, we extracted the recurrent and co-actionable alterations from the molecular signatures of patient cohorts. In each cohort, we first determined the significantly recurrent events from the REFLECT signatures with a permutation-based statistical test (see Methods). The statistical test filters out the features whose observed recurrence is likely by chance in a large patient cohort and preserves the ones that are significantly recurrent, and thus potentially functional. Next, we identified the patient sub-cohorts that carry recurrent and potentially co-actionable events. The co-actionable targets involve (i) the “master biomarker” that defines the cohort and/or (ii) co-occurring actionable biomarkers that are recurrent within a sub-cohort detected by REFLECT. To select actionable mutation and copy number events across sub-cohorts, we matched the oncogenic alterations in REFLECT-signatures to the MDACC and OncoKB precision oncology knowledge bases. To match protein and mRNA expression aberrations to therapeutic agents, we developed a preclinical resource based on the drug target information from MDACC Precision Oncology Knowledge Base, OncoKB, GDSC and CTRP drug-target annotations as well as improvements through literature and expert curation (Suppl. Table S6 and S7)^19,20^. The core assumption in the mRNA/proteomics-based actionable event selection is that overexpression of a drug target predicts potential sensitivity to the respective targeted agent. For example, the increased phosphorylation and total levels of signaling molecules in key oncogenic pathways (e.g., AKT/PI3K/mTOR, RTK, MAPK, cell cycle, DNA repair pathways) are linked to their respective and well-characterized inhibitors (Table S6). Our resulting proteomics and transcriptomics based resource matches 82 proteomic and 89 mRNA aberrations to targeted therapy options.

The REFLECT analysis led to 1,297 and 605 unique drug combinations, of which 374 involve an FDA approved agent, matched to patient sub-cohorts with recurrent features (P-recurrence < 0.05) based on mutation and copy number alterations, respectively. For protein and mRNA expression aberrations (Z-expression > 2 categorized as an aberration, P-recurrence < 0.05 categorized as recurrent), 1,016 and 811 unique combination targets matched to FDA-approved drugs, respectively. With the current Z-expression and P-recurrence thresholds, 31% and 26% of patients with diverse cancers in the meta-cohort are matched to at least one combination therapy involving FDA-approved agents for protein and mRNA analyses, respectively. A total of 2,201 combinations that involved FDA-approved agents were matched to distinct REFLECT signatures based on mutation/copy number, mRNA and protein data. The outcome is a comprehensive precision combination therapy resource that matches drug combinations to patient groups defined by recurrent co-alteration signatures.

### REFLECT-predicted drug combinations can introduce significant therapeutic benefit

We tested whether REFLECT-selected drug combinations have significantly higher therapeutic benefit in the matched molecular backgrounds using independent data sets. First, we devised a validation scheme based on three comprehensive drug combination screens that were also used in the previous DREAM challenges (Figure 3A). We used the Loew’s additivity synergy scores and drug target information for all compounds in the screens as assessed within the DREAM challenge^41^. We identified the subset of the drug combinations in the screens that target the REFLECT nominated co-alterations in the matched cell lines (Figure S9). Next, we compared the therapeutic benefit as quantified by the synergy scores between the REFLECT-nominated drug combinations vs. all other drug combinations (Figure 3B-D, S8). We have demonstrated REFLECT-nominated drug combinations tailored to specific molecular backgrounds have significantly higher synergies compared to other combinations. The therapeutic benefit monotonically increased with increasing recurrence (low P-recurrence) for both mRNA and protein-based predictions and increasing expression levels (high Z-scores). The median synergy scores increased from 7.3 for all drug combinations to 26.1 (p=0.03), 16.2 (p=0.05), and 25.0 (p=0.06) for combinations matched to significantly recurrent (p-recurrence < 0.05) proteomic, mRNA and DNA (mutation/copy number) co-alterations respectively (Table S12). The significant difference in synergy scores is particularly striking given that the REFLECT predictions rely solely on omic backgrounds in a therapy naïve patient cohort but no drug-response data. We conclude REFLECT has the potential to facilitate precision medicine applications that depend on molecular profiling prior to response assessment.

**Figure 3.**
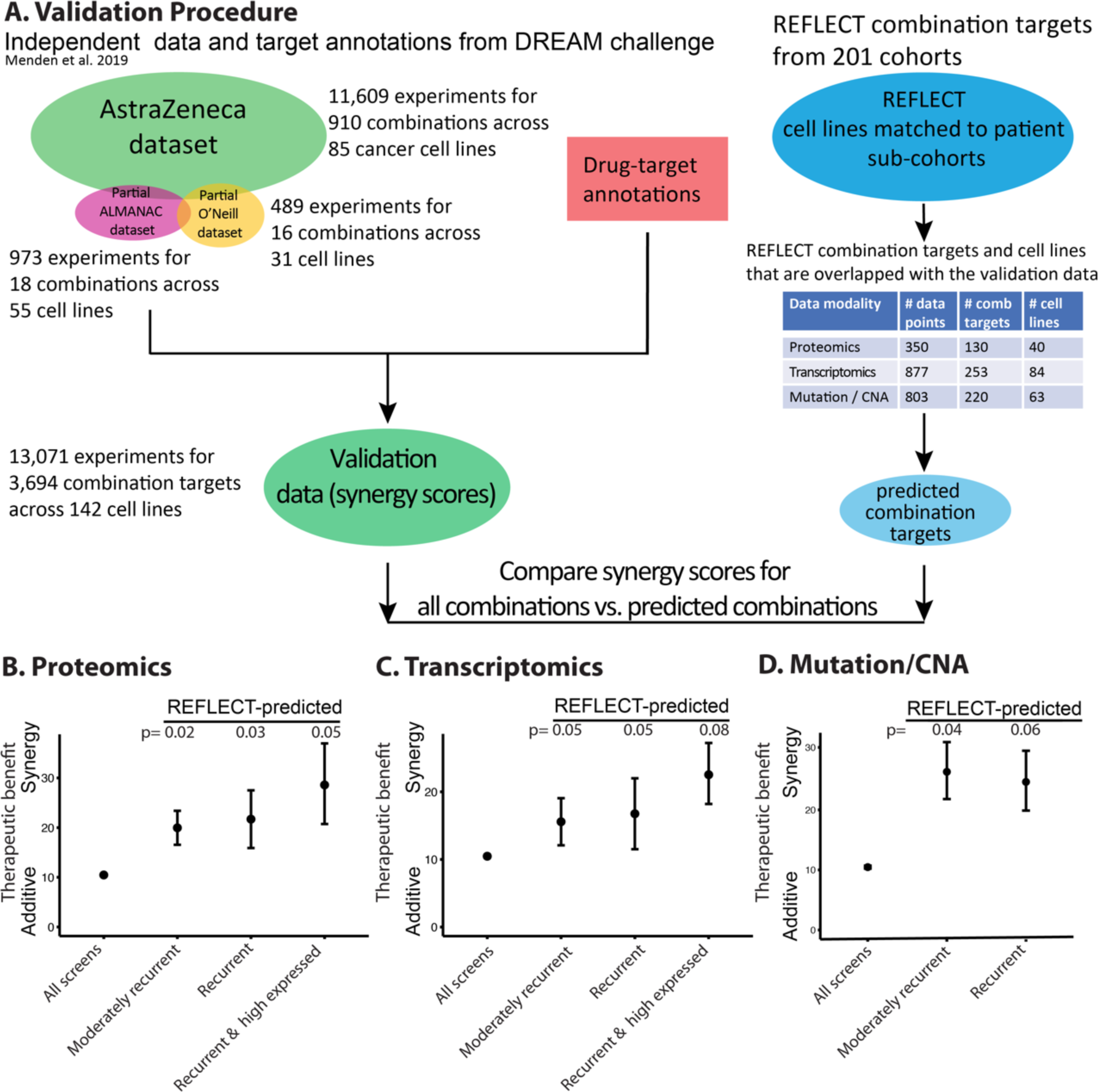
Validation of therapeutic benefit by REFLECT with independent drug response data. A. The REFLECT validation scheme benefits from an independent data set involving the responses to drug combinations based on Astra Zeneca resource, NCI ALMANAC resource and O’Neil et al^39,41^. The drug synergy scores and drug target annotations were integrated within the framework of previous DREAM challenges^39^. To demonstrate the potential therapeutic benefit by REFLECT, we compare the synergy scores of the drug combinations targeting REFLECT selected co-alterations vs. the pool of drug combinations from drug combination screens. The synergy scores for the **B**. proteomics, **C**. mRNA-expression, and **D**. mutation/copy-number driven REFLECT selections are compared. All screens: all readouts from the screns^39-41^. Moderately recurrent: Readouts for REFLECT selections with P-recurrence < 0.25, Recurrent: P-recurrence < 0.05. High expressed: expression Z-score > 2. The therapeutic benefit is quantified by the Loewe’s synergy score (data imported from ref 39). Error bars correspond to standard errors of the mean.

### The immunotherapy response signatures are enriched with inflammatory signals and actionable DNA repair pathway markers

A key PD1/PD-L1 inhibitor response predictor is high PDL1 expression in tumor cells and immune cell infiltration into the tumor microenvironment. To nominate precision combination therapy markers that may improve responses to PD1/PD-L1 inhibitors, we determined the multi-omic signatures and co-occurring actionable targets within a cohort that carry likely responders to immunotherapies (Figure 4). The potential immunotherapy responders are selected based on high PD-L1 levels as well as high tumor infiltrated leukocyte (TIL) fraction. To establish a robust PD-L1 high cohort, we included the tumors whose PD-L1 expressions are in the top quartile across all the pan-cancer meta-cohort for both mRNA and protein levels. Next, to determine tumors with high immune content, we included the samples in the top quartile of TIL fraction across all PD-L1 high tumors. The TIL fraction had been quantified with a mixture model of tumor subpopulations based on methylation probes with the greatest differences between pure leukocyte cells and normal tissue^42^. The resulting high PDL1/TIL cohort, that we call the immunotherapy cohort, contains 406 patients from 23 disease types (Table S4). We then generated the REFLECT-based significantly recurrent mutational, mRNA and phosphoproteomic signatures co-occurring with high PDL1/TIL. An analysis of PD-L1 expression and infiltrating lymphocytes led to a patient cohort with >90% overlap and resulted in virtually identical signatures to those from infiltrating leukocytes.

**Figure 4.**
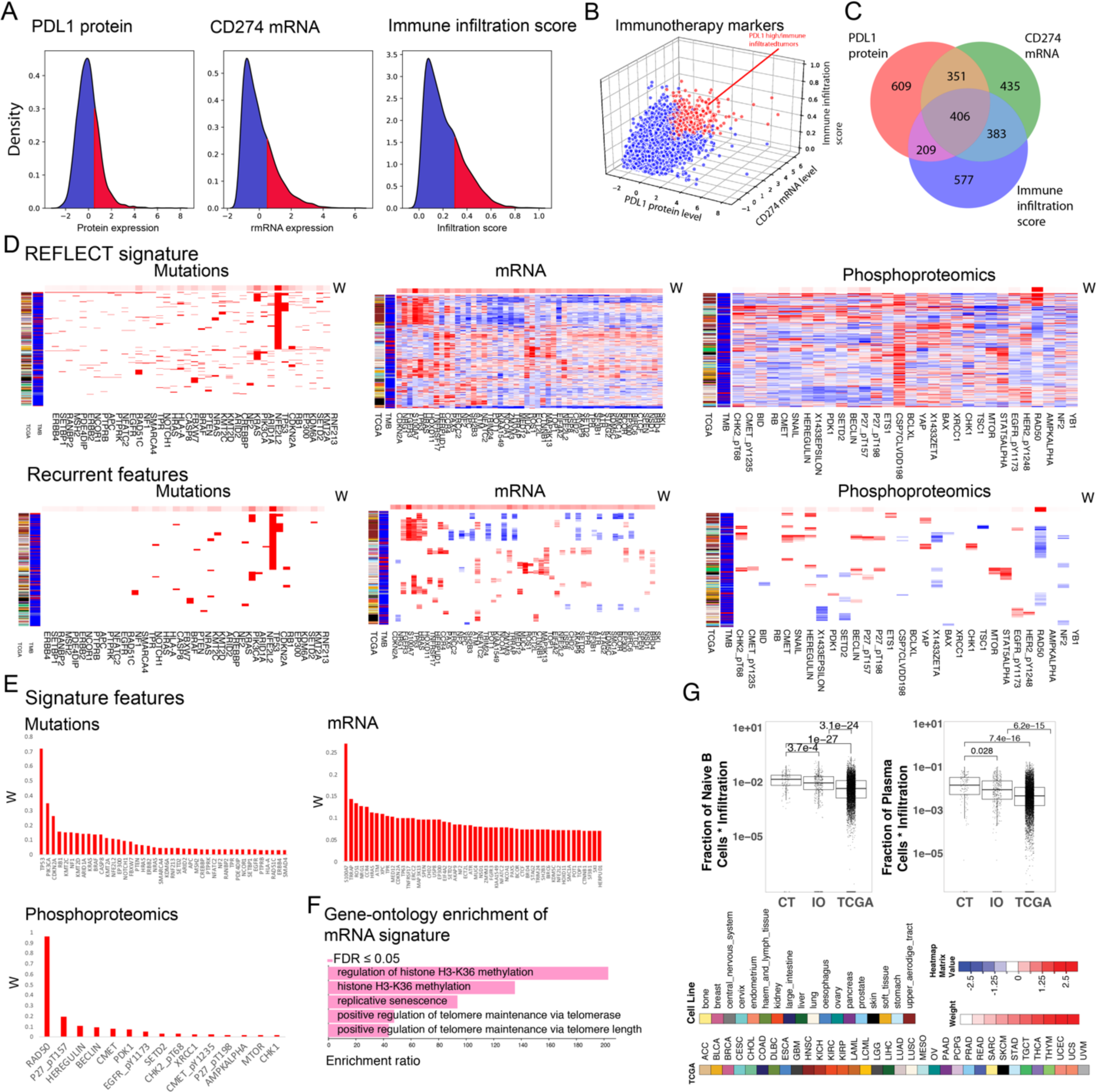
REFLECT analysis of the immunotherapy response marker (IO) cohort. **A-B**. The patients carrying the potential immunotherapy response markers are selected. The patient group that corresponds to the intersection of upper quartiles of PDL1 protein expression, CD274 (gene coding PDL1) mRNA expression and immune infiltration are included. **C**. The intersection of the cases carrying the markers led to a cohort of 406 patients. **D**. The REFLECT signatures (top panel) and significantly recurrent features (bottom panel) for mutations, mRNA expression and phosphoproteomics. **E**. The most discriminant DNA, mRNA and protein features based on REFLECT-analysis in the IO cohort. **F**. The cellular functions that are significantly enriched in the IO mRNA REFLECT signature as detected by functional enrichment analysis. **G**. The statistically significant enrichment of naive B-cells and plasma cells in the CCR4 high/TNFSFR17 (CT) high sub-cohorts within the IO cohort (all IO cases including CCR4 high/TNFSFR17 high sub-cohort).

The mutation analysis of the immunotherapy cohort identified a signature of eight genes (w > 0, P-recurrence < 0.05) covering the commonly mutated oncogenes in cancer (Figure 4D-F). The most recurrent signature genes (P-recurrence < 0.0001) discriminating patient sub-cohorts are the TP53 (50% of the cohort), PIK3CA (15%) and CDKN2A (10%) followed by RB1, BRAF, and KRAS. The mRNA REFLECT signature of the immunotherapy cohort is characterized by a combination of significantly recurrent events (w > 0, P-recurrence < 0.05) in immune-associated genes as well as DNA damage response and repair pathway markers. The strongest contributing mRNA feature is the overexpression of S100A7 (w=0.27, P-recurrence < 0.0001). In the context of cancer, S100A7 protein functions in the tumor microenvironment with roles in inflammation-driven carcinogenesis and recruitment of immune cells including tumor-associated macrophages (TAM) and is highly expressed in a series of cancers including head and neck cancers^21,22,23^. The S100A7 enriched sub-cohorts also carried upregulation of oncogenic genes including overexpression of TP63, HRAS, and FGFR3 particularly in head and neck as well as bladder and squamous lung cancers. Another patient sub-cohort populated by lung squamous cancer cases was enriched with a co-occurring expression of the immune chemotactic receptor CCR4, which is associated with infiltration by diverse lymphocytes and TNFRSF17, a marker of B-cells^24,25^. Indeed, we demonstrated, based on a CIBERSORT analysis, that the tumors in the high CCR4/TNFRSF17 expression sub-cohort are significantly enriched with naive and plasma B-cell markers (Figure 4G).

The transcriptomic signature is also enriched by chromosome organization, cell cycle checkpoint and DNA repair genes such as *TRRAP, CDKN2A, ATM, XPC, ERCC2, CHD2, SETD2, ATR, NSD1, BRD4 and ZYMIM3*. A functional enrichment analysis using the WEB-based GEne SeT AnaLysis Toolkit’ (WebGestalt)^26^ revealed highly significant enrichment of the gene sets involved in histone methylation (FDR-p<0.001) as well as DNA repair associated processes, replicative senescence (FDR-p=0.03) and telomere maintenance (FDR-p=0.01). Similarly, in the phosphoproteomics analysis, depletion of RAD50, a key enzyme for repairing the DNA double-strand breaks, has the highest contribution to discriminate the immunotherapy sub-cohorts with enrichment in two patient sub-cohorts (Figure 4E-F). The first RAD50 depleted sub-cohort (N=111, P-recurrence < 0.0001) is populated by diverse cancer types while the second sub-cohort (N=48, P-recurrence < 0.0001) is enriched by bladder cancers. Increased CHK1 levels (P-recurrence < 0.0001, N=25) and CHK2 phosphorylation at T68 (P-recurrence < 0.0001, N=52), a marker of DNA damage response and checkpoint, also co-occurred within a fraction of the RAD50 depleted sub-cohort, suggesting aberrant DNA repair and damage in such tumors. The enrichment of DNA repair and checkpoint aberrations suggest potential benefit from inhibitors that target individual repair enzymes, particularly the PARP inhibitors that are synthetically lethal with such aberrations. In the context of emerging PARP and PD-L1 inhibitor trials, the sub-cohort specific co-occurrence patterns may guide the selection of patients that are likely to benefit. Such co-occurrence patterns also warrant further preclinical testing to inform novel combination therapy trials that co-target the aberrant DNA repair proteins listed here and immune checkpoints.

### Recurrent co-alterations within DNA repair pathways nominate PARP-inhibitor based combination therapies

PARP inhibitors have provided significant therapeutic benefit particularly for a fraction of cancer patients based on synthetic lethality with DNA repair defects. Here, we have nominated combination precision therapies, in which PARP and other DNA repair inhibitors serve as the anchor compound. We applied REFLECT to patient cohorts marked with alterations (master biomarkers) in 22 DNA repair genes that are implicated in BRCAness and potentially synthetically lethal with PARP (or other DNA repair enzymes) inhibition as listed in the MDACC Precision Oncology Knowledge Base (Table S8). First, we generated the multi-omic signatures for each of the cohorts to identify recurrent co-alterations, and next, selected potential combination therapies using the forward drug selection criterion (Figure S3A and S3C, also see Methods).

To achieve an integrated analysis of the signatures associated with the DNA repair alterations, we mapped the proteomic co-alterations paired with cohort markers on a bipartite graph. On the graph, each recurrent feature within a cohort is linked to the respective co-occurring master marker of the cohort (i.e., the DNA repair aberration that defines a cohort) (Figure 5A-B). We observed an enrichment of aberrant expression in DNA repair proteins (RAD51, ATM, CHK1/2), RTKs (p-EGFR, p-HER2, MET, KIT, VEGFR) and downstream signaling proteins (p-AKT, mTOR, p70S6K, p-JNK, p-SRC, pP38/MAPK14, p-PDK1, p-RAF, p-MAPK) that co-occur with the master biomarkers in the DNA repair family. Guided by the graph enrichment of REFLECT-detected co-alterations in DNA damage markers, RTKs and downstream signaling, we experimentally co-targeted the PARP and co-occurring actionable aberrations (Figure 5D). In drug perturbation experiments, we used the cell lines molecularly matched to (co-cluster with the) specific patient sub-cohorts carrying the co-alterations (Figure 5). We prioritized molecularly matched gynecological and breast cancer patient sub-cohorts and lines as PARP targeting has demonstrated strong but transient therapeutic benefit for a fraction of high grade serous ovarian and basal-like breast cancers. Using the forward drug selection (Figure S3A and S3C) in the RAD50, PALB2 and ATM altered cohorts, we identified sub-cohorts characterized by statistically significant recurrent Chk2 phosphorylation, AKT phosphorylation and MET expression, respectively (P-recurrence < 0.25). For each sub-cohort, we identified a cell line with a matched co-occurrence pattern, namely MFE-296 (RAD50 mutation with recurrent Chk2 phosphorylation), HEC50B (PALB2 mutation with recurrent AKT phosphorylation), and DU4475 (ATM mutation with recurrent MET overexpression). The drug combinations involving PARP inhibitors and the corresponding compounds targeting the recurrent actionable aberrations produce additive to synergistic effects in each of the tests (Figure 5C-E). The experimental tests have provided a proof-of-concept validation that targeting co-alterations may lead to substantial therapeutic benefit to specific patient groups.

**Figure 5.**
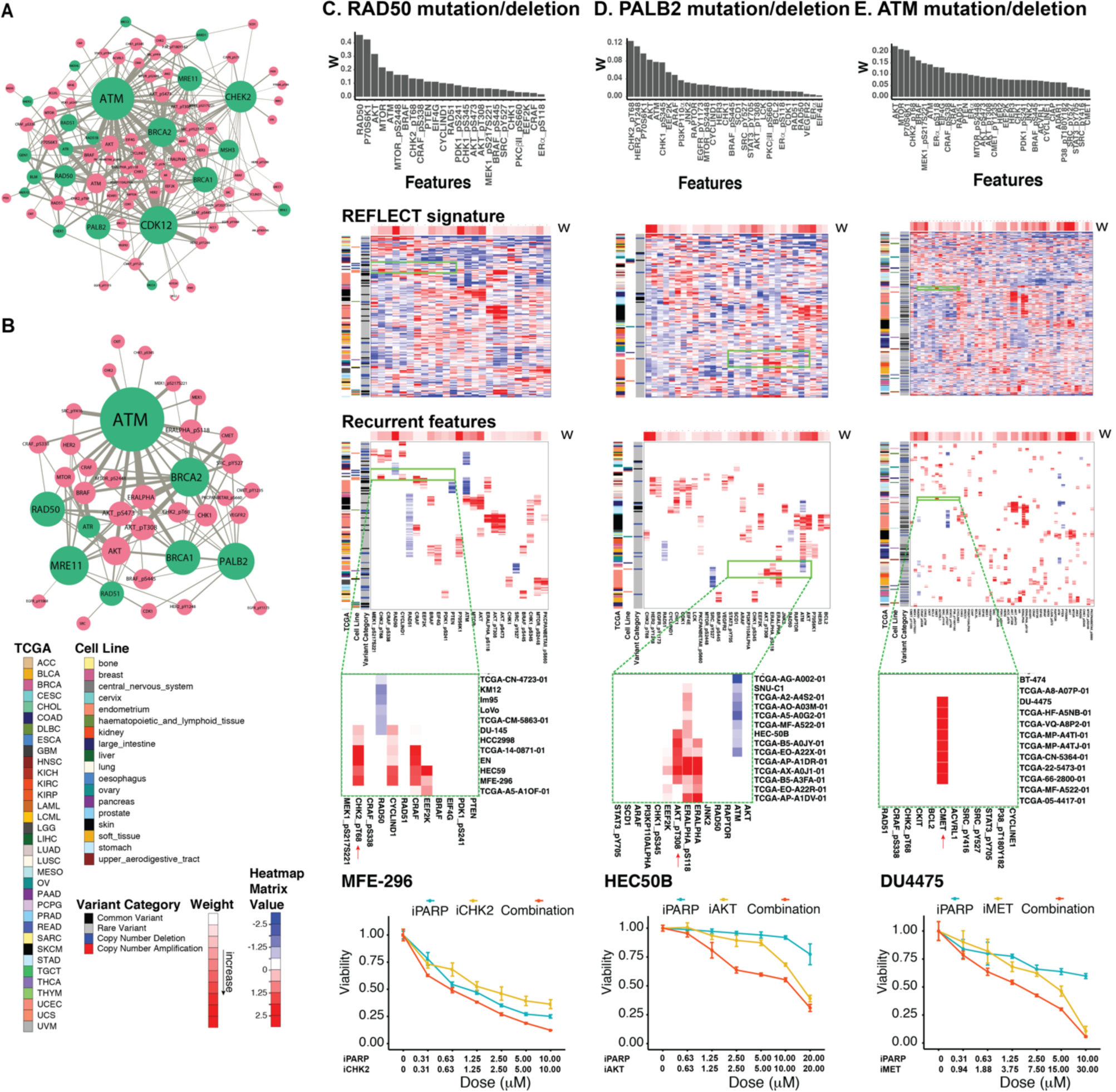
Targeting co-alterations in tumors with DNA repair deficiencies. **A**. The bipartite graph shows the links between the DNA repair genes (master biomarker, green) and co-occurring, actionable protein alterations (REFLECT detected, pink). The size of a node is proportional to the degree of connectivity that represents the number of co-alterations with the master marker. The width of an edge is proportional to the number of patients that carry the co-altered nodes. **B**. The links between ‘core’ repair genes and actionable proteins matched to FDA-approved drugs. **C – E** REFLECT analysis of the mutation/deletions of RAD50, PALB2, and ATM, respectively, which potentially demonstrate synthetic lethality with PARP inhibitors. The top row shows weights of discriminant features from sparse hierarchical clustering, the 2^nd^ row is the REFLECT-signature of each aberration, the 3^rd^ row displays recurrently up/down-regulated protein expressions (Z scores) (p-value < 0.25), the 4^th^ row magnifies a specific region with recurrent patterns and matched cell lines, and the bottom row shows proof-of-principle validations in the matched cell lines. The cell lines are treated with drugs targeting the co-alterations and profiled for cell viability. In the bar charts and heatmaps, only the actionable features with non-zero weights (w) are shown.

### Combination precision therapies for HER2 activated tumors

Therapeutic targeting of HER2 has improved clinical outcomes particularly in HER2+ breast cancers yet resistance to therapy remains a major clinical problem. We used the REFLECT pipeline and a reverse search strategy (Figure S3B and S3D) to identify patient cohorts that may benefit from combination therapies involving HER2 inhibitors (Figure 6). In the reverse search strategy, we scanned proteomic signatures within 201 cohorts, each defined by the actionable biomarkers. We identified patient sub-cohorts carrying actionable oncogenic aberrations (master biomarkers) and increased HER2 total or phosphorylation (p-Y1248) levels that may predict benefit from HER2 inhibitors. The analysis led to the identification of 69 sub-cohorts that carry a HER2 activation marker (P-recurrence < 0.05) alongside the actionable master biomarker. As a result, 338 patients (4% of the TCGA meta-cohort) were matched to 51 unique combinations involving HER2-targeting (Table S10). For proof-of-principle validation experiments, we identified cell lines that map to sub-cohorts of increased HER2 activity. The molecularly matched cell lines were sampled within the cohorts of (i) AURKA mutation/amplification (cell line: BT474), (ii) MAP3K4 mutation/amplification (cell line: SKOV3), (iii) CDKN1B mutation/loss (cell line: HCC1954), (iv) SRC amplification (cell line: SKBR3), (v) PTEN loss/mutation (cell line: MDAMB468). The cell lines were perturbed with HER2/EGFR inhibitors and agents targeting the cohort biomarkers. The drug combinations targeting the co-alterations substantially improved responses over every single agent (Figure 5).

**Figure 6.**
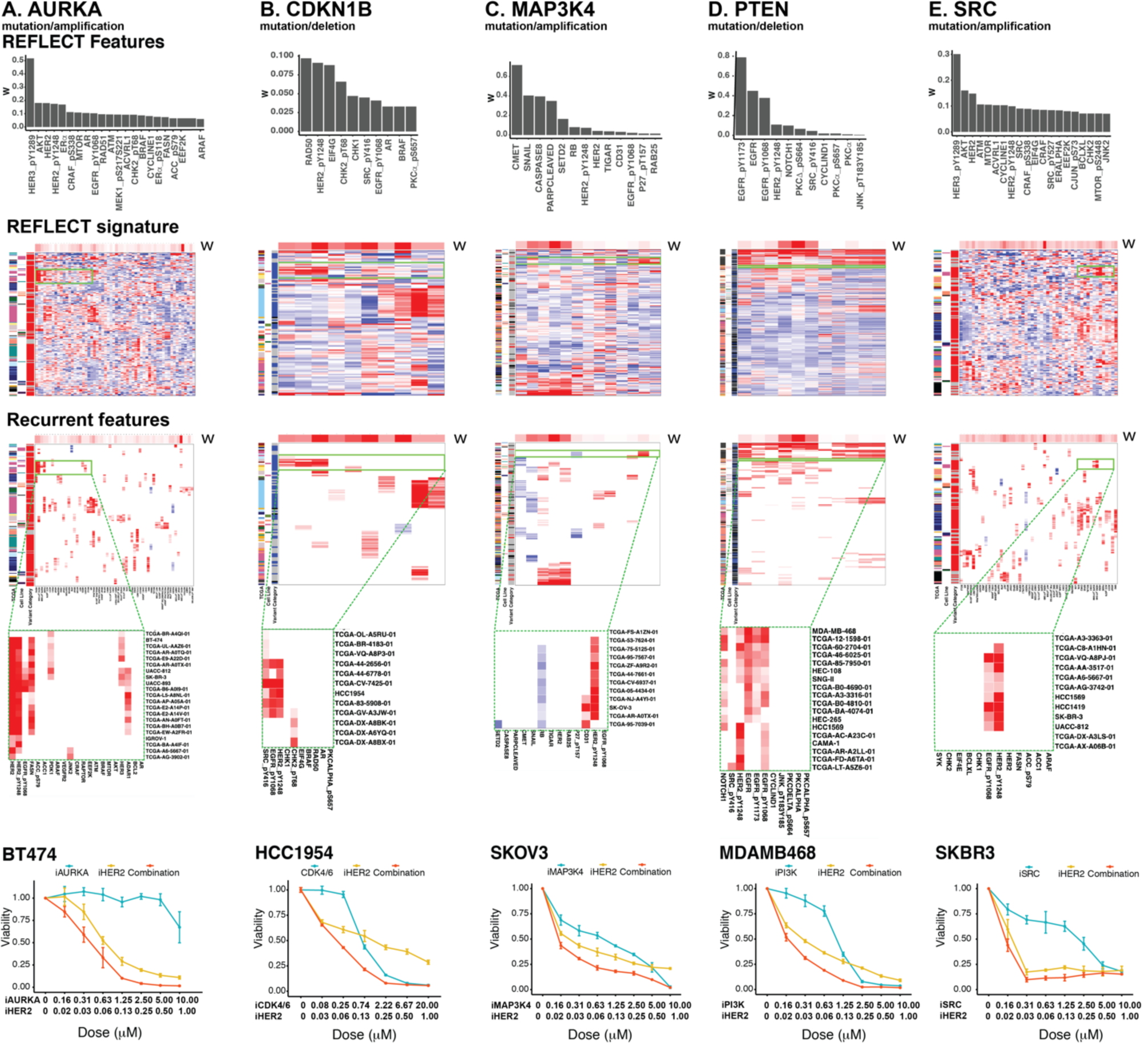
Co-targeting strategies in HER2 activated tumors. Co-targeting of HER2 activation with **A**. AURKA mutations, **B**. CDKN1B loss, **C**. MAP3K4 mutations, **D**. PTEN loss, and **E**. SRC activation. The top row shows weights of features from sparse hierarchical clustering, the second-row displays REFLECT signatures, the third row displays heatmaps of recurrently up/down-regulated protein expressions (Z scores) (p-recurrence < 0.25), the fourth row magnifies the region with the selected recurrent pattern (e.g., HER2/HER2_pY1248 upregulation) and matched cell line. The last row shows proof-of-principle tests in matched cell lines. The cell lines are treated with the combinations of HER2/EGFR inhibitor (neratinib) and inhibitors of AURKA, CDK4, JNK1/2, AKT and SRC. In the bar charts and heatmaps, only the actionable features with non-zero discriminant weights (w) are shown, except for column C (MAP3K4), where all non-zero features are displayed.

## Discussion

Here, we provide a bioinformatic analysis method, REFLECT, to nominate combination therapies tailored to combinatorial aberration signatures in patient cohorts. The REFLECT method includes a machine learning algorithm for the selection of recurrent co-alterations within distinct patient sub-cohorts and benefits from precision medicine knowledge bases to match the co-alterations to therapies. With a REFLECT-based analysis, we have generated a resource that identifies combination therapies matched to > 10,000 patients across > 200 patient cohorts each defined by the existence of an actionable target. REFLECT introduces significant therapeutic benefit as shown with systematic validations across comprehensive drug combination screens. To demonstrate the capabilities of REFLECT, we focused on signatures of three clinically relevant events –potential immunotherapy response markers, DNA repair and damage response aberrations, and HER2 activation. The analysis of HER2 and DNA repair aberrations has led to the determination of patient sub-cohort specific combination therapy nominations that we validated in matched cell lines. The experimental validations revealed either novel drug synergies such as co-targeting Aurora kinase and HER2 in characterized patient groups or established the potentially benefiting patient cohorts for previously tested drug combinations such as those co-targeting PARP with Chk2^27^, AKT^28,29^ or MET^30^ and HER2 with CDK4^31^and PI3K^32^. We have also determined enrichment of specific inflammation and immune cell (B-cells) markers as well as DNA repair alterations within the patient cohorts characterized by high PDL1 expression and immune infiltration, the two biomarkers of immunotherapy response^33^. The co-enrichment of DNA repair/damage response and immunotherapy response markers establishes a rationale for testing PD1/PDL1 inhibitors in combination with DNA checkpoint and repair inhibitors in molecularly matched patiens^34^.

REFLECT is built on a set of core principles in cancer genomics and therapeutics. First, oncogenesis and cancer progression are driven by multiple co-existing driver aberrations^11,35,36^. Such co-existing aberrations may be causally coupled to each other and it is a plausible speculation that targeting such coupled events may lead to significantly synergistic/strong responses. Alternatively, co-alterations may drive parallel and relatively independent events with no trivial causal link in the same or separate tumor sub-clones, yet, may contribute to cancer progression in an additive manner. Targeting the independent events may lead to additive therapeutic benefit with potentially increased durability. Second, we emphasized recurrence in analysis of co-alteration signatures. Oncogenic alterations that are significantly recurrent across patient cohorts are likely functional as such recurrence across samples suggests oncogenic selection to drive oncogenic hallmarks^37^. The significant recurrence of actionable features also suggests the resulting therapy regimes may benefit substantially large patient populations. Finally, precision therapies are usually matched to functional variants of an oncogene (e.g., BRAFV600E). Here, when selecting potentially actionable mutations, we have not filtered out the variants of unknown significance (VUS) and included all variants of a master biomarker if the variant exists in more than 2 patients in the meta-cohort. This approach involves the risk that some passenger mutations that recurred in more than 2 patients are included in the analysis. Yet, we gain the overriding advantage that we detect the REFLECT signatures that are shared between known functional/actionable variants and select VUS for master biomarkers. Such overlaps in the REFLECT signatures between known functional variants and VUS may guide therapeutic options to target potentially important VUS. In summary, the success of our approach relies on the justified observation that recurrent co-alterations that co-activate multiple oncogenic processes may prevent benefit from monotherapies^8^. Yet, such co-alterations create vulnerabilities to matched combination therapies.

The contributions from bioinformatics tools have become more needed as multi-omic data-driven clinical studies such as the WINTHER trial are emerging^38^. Proteomics-based markers that indicate activation of actionable oncogenic pathways can also complement genomics-based patient selection and drug sensitivity prediction. The proteomic modality is less explored and provides a major opportunity as most targeted agents impact protein activities to mediate the therapeutic effects. For example, increased EGFR and HER2 activating phosphorylation levels is a predictive marker for the EGFR/HER2 inhibitors while nearly half of HER2/EGFR-mutated tumors do not lead to high levels of EGFR/HER2 activating phosphorylation. Proteomic changes, however, may not serve as the sole criterion for drug selection, at least due to the current regulatory limitations around the use of proteomics for clinical tests but is an important complement to other data types. By assessing multiple data modalities at once, we expect tools such as REFLECT will improve the fraction of benefiting patients while providing more reliable target matching in preclinical and data-guided clinical trials. In this respect, the stratification into patient cohorts based on actionable biomarkers emulates the ongoing basket trials such as NCI-MATCH with the added value that it enables matching combination therapies to sub-cohorts with shared molecular compositions.

REFLECT is an informatics platform for therapy selection guided by multi-omics profiling in preclinical and clinical trials. As lack of durable responses to genomically matched monotherapies significantly limits the population of benefiting cancer patients, REFLECT is potentially a critical addition to the current precision medicine paradigm. With expanding data from precision therapy trials and genomics studies, it will become increasingly feasible to match combination therapies to complex molecular signatures. We expect REFLECT will be a commonly used tool and resource to select precision combination therapies in both preclinical and clinical studies.

## Acknowledgment

We thank Funda Meric-Bernstam, Scott Woodman, Gordon Mills, Zeynep Dereli, Chris Wakefield and Murat Cokol for invaluable comments. This work is supported by grants from MDACC Support Grant P30 CA016672 (the Bioinformatics Shared Resource), OCRF Collaborative Research Award, ICI Fund, CPRIT High-Impact/High-Risk Award (RP170640) and funding from MD Anderson Colorectal Moon Shot program.

## Supplementary Methods

### Multi-omic data

Phospho-proteomics data from TCGA resource and cancer cell lines are available from The Cancer Proteome Atlas Portal (https://tcpaportal.org/). We obtained level 4 data (batch normalized across disease types) for both patient cohorts and cell lines. After filtering out samples having missing values, the TCGA patient dataset had 7544 samples from 32 cancer types (no RPPA data is available for LAML cancer type), and the cell line dataset had 397 samples from 18 lineages (MCLP). Therefore, each dataset covered a large number of samples from diverse cancer types. Because of their overlapping coverages of diverse disease types, we assumed that the TCGA patient dataset and the cell line dataset follow the same distribution. To remove the batch effect and merge them together, we standardized each dataset using Z-score transformation so that each has zero mean and unit variance. Then, the two standardized datasets were combined to form an integrated phospho-proteomics dataset. For mRNA expression data, we integrated TCGA patient (N = 8,887) and CCLE cell line (N = 1,156) RNA-seq data. Similar to the integration strategy used for the phospho-proteomics data, we Z-score standardized the log2-transformed mRNA expressions in each dataset and then combined them together. The mutation and copy number data for patient cohorts and cell lines were from TCGA (N = 8,764) and the COSMIC-CLP (N = 996), respectively. For the mutation data, we filtered out hypermutated samples. Then, mutations were discretized to binary values (mutated=1, wide-type=0). Copy numbers were discretized to {-1, 0, 1}, representing deletion, no alteration, and amplification, respectively. Without the need of correcting batch effects for discretized mutations and copy numbers, TCGA patient data and cell line data are combined. For each data modality (phospho-proteomics, mRNA expressions, mutations, copy numbers), the distributions of samples across cancer types or lineages are shown in Figure S1.

### Genomic stratification markers

We selected actionable genomic alterations as the stratification markers from two sources. First, we selected 15 genomic vulnerability markers from NCI-MATCH (Molecular Analysis for Therapy Choice) (Table S2). Then, from 207 potentially actionable alterations from the MD Anderson Precision Oncology Decision Support (PODS), we extracted 182 genomic alteration markers that were not overlapped with the NCI-MATCH markers (Table S3). For each marker, the alterations include mutations, gene amplification/deletions, and specified amino acid substitutions (e.g., BRAF V600 mutation). Depending on specific genes, activating mutations are combined with amplification, and inactivating mutations are combined with gene deletions. These markers also cover the actionable genes listed in OncoKB. Finally, we constructed a cohort of patients carrying potential immunotherapy response markers and 3 other cohorts based on the existence of key oncogenic alterations – MYC amplifications, TP53 hotspot mutations, and KRAS mutations (Table S4).

We termed these markers as “master” biomarkers (N = 201) and listed them in Table S5. Annotations based on Gene Ontology (GO) Biological Process terms were used to categorize the master biomarkers. The top three GO terms were used; terms were ranked by evidence types and counts as provided by GO database. Code is provided at https://github.com/cannin/extract_go_slim_annotation

### Sparse hierarchical clustering

Sparse hierarchical clustering identifies clusters driven by the most discriminant features as follows^12^. The input to the algorithm is X, an *n* ⨯ *p* data matrix with *n* observations (e.g., patients) and *p* features (e.g., genes). *x*_*i*_ is an observation that is a *p*-dimensional vector indexed by *i*, where *1* ≤*i* ≤ *n*, and *d* (*x*_*i*_, *x*_*i*′_) is a measure of the distance between observations *x*_*i*_ and *x*_*i*′_. We assume that the distance measure is additive in features, that is, 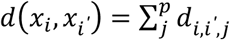, where *d*_*i,i*′,*j*_ is the distance measure between observations *x*_*i*_ and *x*_*i*′_ along feature *j*.

The distance is defined as either the squared Euclidean distance 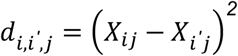 or the absolute difference 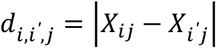. The quadratic Euclidean distance, due to the square operation, assigns a relatively higher weight to larger values, while the absolute difference treats all values uniformly. In REFLECT, to smoothen small differences, the squared Euclidean distance is employed as the distance measure for the continuous data (i.e., RPPA and mRNA expression). For the discrete data (i.e., mutation and copy number alterations), only a small finite set of values exists, that is, {-1, 0, 1}, so we use the absolute difference as their distance measures.

The sparse hierarchical formulation is derived from the standard hierarchical clustering equations. In the standard hierarchical clustering, we solve the constrained maximization problem

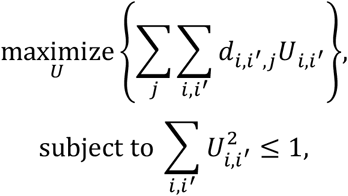

which leads to the optimal solution 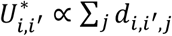.

The iterative sparse hierarchical clustering is formulated as the solution to a weighted constrained optimization problem

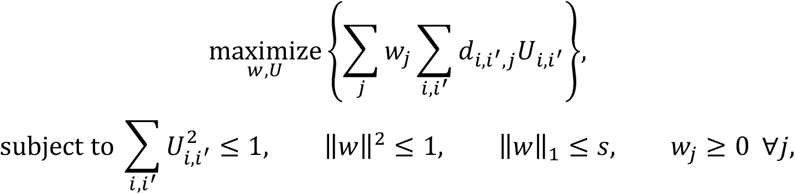

where *w*_*j*_ is a weight for feature *j*, and *s* is a tuning parameter that controls sparsity of *w*. Let *U*^∗∗^ optimize the above criterion, from which we have 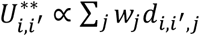, the distance along each feature is weighted according to *w*_*j*_. And due to the constraints, a small value of *s* makes *w* sparse, so *U*^∗∗^ depends only on a small subset of features. Once the w is optimized, hierarchical clustering is performed on *U*^∗∗^. The operation is iterated until convergence (w is fixed; U** is optimized and next U** is fixed; w is optimized), w and hence the sparse clustering. The output is a sparse clustering of observations defined by the distribution of a small set of most discriminant features. In REFLECT, we employ the sparcl R package (implemented by Witten and Tibshirani) to execute the sparse hierarchical clustering.

### Selection of tuning parameter via the Gap statistic

In the sparse hierarchical clustering, the tuning parameter *s*, which controls sparsity of *w*, needs to be determined prior to clustering. We apply a permutation approach called the gap statistic to select an appropriate value for *s*. For a specific tuning parameter *s*, the gap statistic measures the strength of the clustering on real data compared with that expected on a reference null distribution. The computational implementation is as follows.

Step 1: Generate *B* reference datasets {*X*_*1*_, …, *X*_*B*_} by permuting the observed dataset *X* independently within each feature.

Step 2: For each candidate of *s*, where 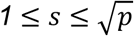:

i. Obtain the optimal objective *O*(*s*) *=* ∑_*j*_ *w*_*j*_ ∑_*i,i*′_ *d*_*i,i*′,*j*_*U*_*i,i*′_ by performing the sparse hierarchical clustering on the data *X*.
ii. For *b = 1*, …, *B*, obtain the optimal objective *O*_*b*_(*s*) by performing the sparse hierarchical clustering on the data *X*_*b*_.
iii. Compute the gap statistic, the deviation between log objective function of clustering with actual data and mean of the log objective function from the null distribution 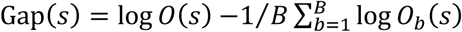.

Step 3: Use the following optimization algorithm to select the optimal *s*^∗^ that maximizes *Gap*(*s*):

i. Search for an *s*_*min*_ such that when *s* ≥ *s*_*min*_, the number of non-zero features from sparse clustering is greater than *N*_*cutoff*_. Here, we choose *N*_*cutoff*_ *= 10* to prevent the selection of an extremely sparse set of features, which may not represent the complexity of the dataset.
ii. Find *s*_*max*_ that corresponds to the maximum gap statistics.
iii. Calculate the mean *E*[*Gap*(*s*)], the standard deviation *SD*[*Gap*(*s*)], and the standard error of the mean *SEM*[*Gap*(*s*)] for all *s*values in [*s*_*min*_, *s*_*max*_].
iv. Find 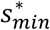 such that 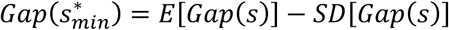.
v. Calculate the mean *EB*[*Gap*(*s*′)] for all *s*′ in 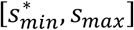. If *EB*[*Gap*(*s*′,*z* > *E*[*Gap*(*s*)] − *SEM*[*Gap*(*s*)], let 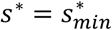; otherwise, let 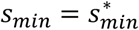 and return to the step (iii).

Using this procedure, we selected tuning parameters for 201 cohorts while performing sparse hierarchical clustering, which determined the number of nonzero features. The distribution of number of nonzero features are shown in Figure S2.

### Robustness of feature selection

We evaluated the robustness of feature selection in REFLECT analysis with a subsampling approach. We quantified the stability of the top features calculated with partial and varying datasets compared to the original dataset. Since this calculation is computationally costly, we performed this analysis selectively for several datasets from cohorts of DNA repair response aberrations and HER2 activation, including RAD50 mutations/deletions, PALB2 mutations/deletions, ATM mutations/deletions, AURKA mutations/amplifications, CDKN1B mutations/deletions, MAP3K4 mutations/amplifications, PTEN mutations/deletions, and SRC mutations/amplifications.

For each dataset, REFLECT was used to get an ordered list of the top *N* discriminant features *L*_*whole*_ =[*F*_(*1*)_, …, *F*_(*N*_)], where *N = min*(*20*, |*w*|_*w*>0_), and |*w*|_*w*>0_ is cardinality of nonzero features. Then, we randomly subsampled the dataset to a specified fraction, *f*, where *0*.*6* ≤*f* ≤*0*.*95* in 0.05 increments, for *S* times, where *S = 200*. The sparse hierarchical clustering was performed on each subsampled dataset to extract an ordered list of top *N*^∗^ discriminant features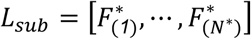, where we chose *N*^∗^ *= min*(*10*, |*w*|_*w*>0_). A score can be calculated as follows by comparing features in *L*_*sub*_ that recapture the ones in *L*_*whole*_:

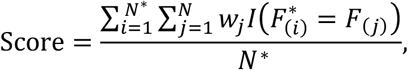

where *I*(·) is an indicator function, *w*_*j*_ represents a rank scaled weight of feature *j* for *1* ≤*j* ≤*N*. The rank-scaling is formulated as

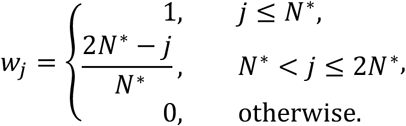

where the top-ranked *N*^∗^ features contribute most to the score while the intermediate ranked *N* − *N*^∗^ features have contributions that are scaled-down in proportion to their ranks. In this scheme a high-weight, high-ranked feature has a higher contribution to the robustness score compared to a low-weight, low-ranked feature. Finally, the robustness scores were visualized using box plots, showing the distribution of *Score* over *B* subsampled datasets as a function of the subsampling fraction *f*.

### Statistical significance of recurrence

In order to assess a p-value for recurrence, we need to calculate the probability of having at least k consecutive altered samples *P*_*val*_(*N, k*), from a total of N samples. A simple way to calculate this probability is by calculating its complement, *P*_*val*_(*N, k*), the probability of having less than k consecutive samples. Here we derive the equation: The p-value of *k* consecutive altered samples by chance can be calculated in the form

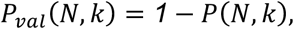

where *P*(*N, k*) is the probability that we do NOT have at least *k* consecutive altered samples.

*N* is the total number of samples, and *N*_*a*_ is the total number of altered samples, where *N*_*q*_ < *N*.

Here, we derive an approximate formula for the calculation of *P*(*N, k*) by assuming the probability of the alteration *p* is a constant, which holds true when *k* is much smaller than the number of altered samples *N*_*q*_. Then, the probability, *p*, of observing an alteration in a sample is calculated empirically by

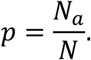

The probability of a sample being unaltered, denoted as *q*, is

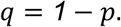

Given a sequence of samples that do not have *k* consecutive altered samples, the end of a sequence must be an unaltered sample followed by *i* altered samples, where *0* ≤ *i* < *k*. Let *P*(*n, k, i*) denote the probability that a sequence of length *n* has less than *k* consecutive altered samples AND ends with *i* consecutive altered samples. Thus, we have

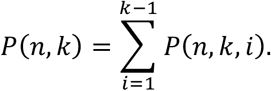

Notice that if we remove *i* samples from one of the borders of *i* consecutive altered samples in the class (*n, k, i*), the sequence belongs to the class (*n* − *i, k, 0*). Thus, we have

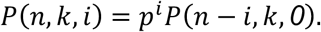

Moreover, if we add one unaltered sample to the end of *i* consecutive altered samples in the class (*n, k, i*), the sequence is in the class (*n* + *1, k, 0*), which gives us the relationship

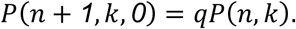

By combing the above three equations, we have

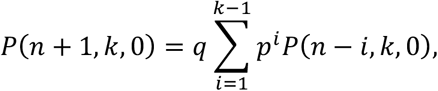

which gives us a linear iterative relationship to calculate *P*(*n, k, 0*). The initial condition is *P*(*n, k*) *= 1* if *n* < *k*. Finally, we can get *P*(*N, k*) expressed in the form

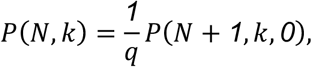

which allows us to get the p-value using *P*_recuurence_(*N, k*) *= 1* − *P*(*N, k*).

### Development of mRNA and RPPA based actionable lists

To get a comprehensive list of actionable targets and their inhibitors, we combined drug-target annotations from 4 databases: (i) MDACC Precision Oncology Knowledge Base (Kurnit, 2018, Table 1), (ii) Precision Oncology Knowledge Base OncoKB (downloaded on Feb, 2020), (iii) Genomics of Drug Sensitivity in Cancer (GDSC), and (iv) Cancer Therapeutics Response Portal v2 (CTRP v2). The combined list includes actionable targets that can be inhibited by FDA-approved drugs, drugs used in clinical trials, and preclinical tests. We defined the mRNA targets based on the assumption that mRNA overexpression of the target predicts sensitivity to the respective agent. Similarly, we obtained proteomics-based actionable targets from the RPPA-assayed proteins for both total protein expression and activating phosphorylation levels. For example, high levels of activating AKT phosphorylation is a target for AKT inhibitors. Finally, we manually curated the generated list based on literature and expert knowledge. Overall, 82 proteomics and 89 transcriptomics based actionable targets were generated (Table S6 and S7).

### Systematic assessment of therapeutic benefits by REFLECT

To evaluate the ability of REFLECT to nominate effective combination therapies, we analyzed the therapeutic effects of REFLECT-selected drug combinations included in comprehensive drug combination screen datasets. The drug combination screen datasets were obtained from supplemental materials of ref 39, including: (i) AstraZeneca dataset (11,706 experiments for 912 combinations across 86 cancer cell lines), which was also used in DREAM challenges^39^(ii) a fraction of the ALMANAC dataset (973 experiments for 8 combinations across 55 cell lines) containing targeted agent combinations^40^ (iii) a fraction of the screen generated by O’Neil et al^41^ (referred as “O’Neil data”) (913 experiments for 10 combinations across 31 cell lines). We combined the data from the three drug screens leading to 13,592 experiments for 926 combination drugs across 143 cell lines. Then the drugs in the data was mapped to their corresponding targets using the annotation provided in supplemental materials in ref 39, and drugs without specific targets were removed. It leads to a dataset of 75,612 unique data points for 3,694 combination targets across 142 cell lines. The combined dataset (referred as the “validation drug screen set”) is used to assess the therapeutic benefits of REFLECT-selected drug combinations tailored to co-occurring molecular aberrations over other combinations. We extracted all the combination targets across cell lines from the 201 REFLECT cohorts that overlapped with the validation drug screen set. In the REFLECT-analysis, there are 350 data points for 130 combinations across 40 cell lines for proteomics, 877 data points for 253 combinations across 84 cell lines for mRNA expressions, 732 data points for 186 combinations across 63 cell lines for mutations, 170 data points for 74 combinations across 23 cell lines for copy number alterations (Table S13). For each data modality, we compared the therapeutic effects of the combinations tailored to REFLECT signatures with the validation drug screen dataset. The therapeutic benefit is quantified by the Loewe synergy scores (imported from ref 39) for each drug combination test. We assessed the therapeutic effects of REFLECT-selected combinations that match to varying recurrence levels (P-recurrence), protein/mRNA expression levels (Z score), and discriminant power (w). For each test, the significance of the REFLECT-based benefits was assessed with P-values calculated against a null-distribution that is generated by bootstrapping from the combined validation set (10,000X, group size=number of REFLECT predictions).

### Selection of nominated combination targets for experimental validation

In REFLECT, the primary target is master biomarker, and the secondary target comes from one of recurrent features identified during clustering. To validate the performance of this computational framework, we performed *in vitro* experimental tests on some of the combination targets nominated from REFLECT analyses.

We employed two complementary selection strategies for validation experiments of combination targets X + Y, where X represents the primary target from one of the genomic stratification markers, and Y is the secondary target from a recurrent (phospho-)proteomic feature. The first strategy is called a forward selection, in which X is fixed, and then Y is selected to be one of the recurrent REFLECT-signatures. On the other hand, in a reverse selection strategy, X is varied while the REFLECT-signature Y is fixed.

We designed two series of validation experiments corresponding to both forward and reserve selection strategies. For the forward selection experiments, we tested combination targets having the primary targets corresponding to master biomarkers that suggest PARP vulnerability (Table S9). These genomic alterations are mutations or deletions of “BRCAness” genes (Table S8). The secondary targets are recurrent features after a REFLECT analysis of each dataset based on those stratified genomic markers. For the reverse selection experiments, we used recurrently upregulated HER2/HER2_pY1248 as the secondary target (Table S10). We ran REFLECT analyses on 201 cohorts and selected primary targets as master biomarkers that co-occur with recurrent HER2/HER2_pY1248 overexpression.

### Cell lines and reagents

The cell lines used in experiments included BT-474, HCC1954, MDA-MB-468, SK-BR-3, and SK-OV-3. They were obtained from MD Anderson Characterized Core Cell Line Facility and were authenticated by fingerprinting using short tandem repeat testing. The absence of mycoplasma contamination was also verified. Cells were maintained in RPMI-1640 supplemented with 10% heat-inactivated FBS and 1% penicillin with streptomycin. Drugs used in the experiments are listed in Table S11.

### Cell culture, growth inhibition, and combinatorial inhibitor effect analysis

Depending on the cell line doubling times, 2000-5000 cells were seeded into a 96-well plate 24 hours before treatment. The cell lines were plated and treated with two individual inhibitors and their combination, in triplicates, over six twofold dilutions. The cell lines were treated in triplicates for different concentrations of the inhibitors from 0 to 10 or 20*μ*M in 6 two-fold dilutions (in 5% FBS) to generate a dose-response curve. The selected doses of each drug were added using Tecan D300e Digital Dispenser. Viability was assessed 72 hours post-treatment using PrestoBlue cell viability reagent (Thermo Fisher Scientific) by measuring fluorescence on BioTek Synergy H1 Hybrid Multi-Mode Reader according to the manufacturer’s recommendation. Background values from empty wells were subtracted and the data was normalized to untreated control. The log of drug concentration was plotted against the growth inhibition (Figure 5, 6, S6).

